# From microbial diversity to function; evaluating dimensionality reduction methods

**DOI:** 10.1101/2025.10.30.685628

**Authors:** Emelia J Chamberlain, William Boulton, Elizabeth J Connors, Theodore Calianos, Jeff Bowman, Jessie M Creamean, Thomas Mock, Heather H Kim

## Abstract

Artificial Intelligence (AI), and more specifically Machine Learning (ML), have become an increasingly prevalent tool in microbial oceanography. The high dimensionality of microbial diversity data from ‘omics observations is highly suitable for ML analysis, with many recent studies showcasing their utility for exploratory ecological feature finding and process prediction. Here, we apply three well-documented dimensionality reduction methods including Principal Coordinate Analysis (PCoA), Self Organizing Maps (SOM), and Weighted Gene Correlation Network Analysis (WGCNA), to near daily 16S rRNA gene amplicon sequencing data from the 2019-2020 MOSAiC International Arctic Drift Expedition. We compare the k-means clustering outputs from these methods to extract functionally distinct seasonal microbial ecotypes in the surface Arctic Ocean. Our results indicate the SOM method outperforms a more traditional PCoA ordination, identifying a greater number of metabolically distinct functional groups. We then investigate the importance of including biological context in dimensionality reduction by comparing functional outputs to a taxa clustering approach using a k-means adapted WGCNA correlation network. Regardless of data input, all 3 methods identified 3-4 recurrent ecotypes with distinct taxonomic and functional cut-offs driven by seasonality, water mass, and substrate turnover. Ultimately, these results reinforce such methodologies as a meaningful translator in the mining of historical amplicon datasets to address modern mechanistic questions and incorporate greater ecotype diversity into mechanistic biogeochemical models.

**Importance:** Connecting microbial community structure to ecosystem function is an important step in accurately modeling climate-relevant biogeochemical processes yet remains a major challenge in microbial oceanography. This manuscript demonstrates how emerging machine learning approaches can establish this connection by uncovering recurrent ecological patterns in Arctic Ocean microbial communities. Using near-daily 16S rRNA gene and supplementary metagenome data from the MOSAiC drift expedition, we identified distinct “ecotypes,” or groups of microbes that perform differentiable functional roles within the ecosystem. Importantly, our methods reveal new connections between microbial identity and function that traditional analyses may overlook. It is possible such techniques could be applied to historical amplicon datasets, allowing scientists to revisit and reinterpret existing data to better understand how polar ecosystems are responding to environmental change and to improve future predictive climate models.

## Introduction

Microorganisms represent the most abundant and diverse form of life on Earth (1). The tremendous taxonomic and functional diversity of microbial communities serves as a fundamental driver of ecosystem functions, marine carbon and biogeochemical cycles, and trophic energy transfer, with different species and functional groups contributing uniquely to these processes (2). Among these microbes, marine heterotrophic bacteria (hereafter bacteria) and archaea are particularly important, processing vast quantities of carbon through their complex metabolic networks. Bacteria alone respire up to 85-90% of the 50 petagrams of carbon fixed annually by phytoplankton (3, 4), an amount equivalent to half of global photosynthesis, making them essential players in regulating whether this carbon returns to the upper ocean-atmosphere boundary as CO_2_ or sinks to the deep ocean for centuries to millennia (5–7). However, the role diversity plays in regulating this process is not well constrained and certain bacterial traits, like maximum respiration rate, can vary widely among microbial groups (8), with the most abundant taxa contributing little to overall rates (9).

Over recent decades, advances in sequencing technologies have enhanced our ability to profile bacterial communities, yielding vast, ‘omics datasets (10). As the importance of microbial contributions to climate processes becomes increasingly recognized, so too does the need to translate these complex community patterns into functional units that can inform predictive biogeochemical models (11). Machine learning (ML) has emerged as a powerful approach for reducing dimensionality across complex microbial datasets and identifying ecologically and environmentally relevant features. While all ‘omics data are rich in biological information, high-dimensional amplicon sequence variant (ASV) data are particularly suitable for ML techniques (12). Recent studies have highlighted this, using both supervised and unsupervised ML methodologies to distill the complexity of microbial community structure into trait-based groupings, i.e., “ecotypes” linked to specific ecosystem functions (13) and extract the best taxonomic predictors for key biogeochemical rate processes such as bacterial production (14) and oxygen utilization (15).

Highlighting the field’s excitement around these methods, a flurry of recent reviews have further expanded upon why and how to apply this growing toolkit for microbial ecologists. Qu et al. (2019) (16) demonstrated how supervised classifiers can predict host phenotypes from microbiome profiles, highlighting the potential of ML to capture complex ecological patterns. Topçuoğlu et al. (2020) (17) introduced an interpretable and reproducible ML pipeline tailored to microbiome data, addressing barriers around accessibility and transparency in ecological applications. Bowman (2021) (18) highlighted the predictive capacity of microbial ML applications, while Hernández Medina et al. (2022) (19) emphasized integrating environmental metadata and outlined modeling-specific considerations for microbiome data. Most recently, Asnicar et al. (2023) (20) led a discussion on the basic tenets of ML for microbiologists and challenges related to interpretability and standardization. While ML shows clear utility for microbial trait extraction, mechanistic connections between microbial community composition and ecosystem-level processes remain underexplored, despite evidence suggesting their value for ecosystem modeling (8). However, the translation of microbial community-level patterns into functional representations suitable for ocean biogeochemical models presents significant methodological and conceptual challenges. As computational approaches increasingly interface with these predictive ocean models, understanding which ML frameworks can effectively translate microbial community composition into the functional units needed for such modeling frameworks becomes increasingly relevant.

In this study, we compare and evaluate three dimensionality reduction approaches compatible with k-means clustering as a strategy to navigate the high-dimensionality challenge of bacterial data: (1) Principal Coordinate Analysis (PCoA) (21), (2) Self-Organizing Maps (SOMs) (22), and (3) Weighted Gene Correlation Network Analysis (WGCNA) (23). While all three approaches preserve the underlying structure embedded in high-dimensional microbial datasets, they differ significantly in assumptions, outputs, and interpretability. Each method offers distinct advantages: PCoA provides traditional statistical ordination that reduces compositional data noise by projecting it into a lower-dimensional space (24). SOMs employ fully unsupervised, nonlinear neural networks using a competitive learning algorithm, making them robust tools for data exploration and visualization (25). WGCNA constructs weighted correlation networks to identify discrete modules of co-varying taxa, offering a more taxa-driven mechanistic framework suitable for hypothesis-driven ecological analysis (23).

We test these approaches using a multi-seasonal bacterial dataset from the surface waters of the Arctic Ocean, in a region where climate warming is occurring at almost four times the global average rate (26). This ocean represents an ideal testbed for comparing dimensionality reduction methods due to its extreme seasonal bacterial shifts (27) and the critical roles these communities play in carbon cycling processes that could significantly impact Arctic amplification (28). The rapid environmental transitions in the Arctic have the potential to release previously sequestered carbon (29), while fundamentally altering bacterial community structure and function (30). Identifying which ML methods best capture these complex ecological patterns in high-dimensional Arctic bacterial datasets is therefore essential for improving model predictions of regional and global climate responses mediated by microbial feedbacks.

## Materials and Methods

### Sample Selection and Collection

The Multidisciplinary drifting Observatory for the Study of Arctic Climate Expedition (MOSAiC; 2019–2020) was an interdisciplinary year-long drift experiment in the central Arctic Ocean that created a wealth of ecological (31) and genomic (10) data which we have leveraged for this study. From the publicly available MOSAiC 16S rRNA gene amplicon sequencing dataset (32), a subset of 215 suitable surface seawater samples were selected and assessed. These were collected as close to daily as possible (average sampling time 15:00 UTC) between 10/29/2019 and 09/18/2020 over the course of the MOSAiC drift, covering a 10º latitudinal gradient 89.1º – 79.1º across the Amundsen and Nansen Basins (31). Data gaps exist where project personnel were not on board (12/13/2019 – 02/20/2020) and during periods of ship relocation (07/31/2020 – 08/22/2020). Samples were collected from the underway seawater system (RV Polarstern; 11m inlet), with highly comparable community structure results to samples from similar depths collected using a CTD Rosette (15). Additionally, 18 of the underway sampling dates matched with whole metagenome data collected at either 10 or 11 m depth from the CTD rosette (10). Ancillary environmental data (surface temperature and salinity) were collected continuously from the underway system and corrected as described in (33). Chlorophyll-a concentrations were collected and filtered at the same time as collections for microbial community structure and are described by Hoppe et al. (2023) (34).

### DNA sequencing and metabolic function

Sample and data processing for water column bacterial and archaeal community structure (16S rRNA gene amplicon sequencing) is described in detail by Chamberlain et al. (2025) (15). Briefly, 1 L of underway seawater was filtered through a 25mm 0.2µm Pall Corporation Supor membrane filter and frozen (–80ºC). DNA was extracted using a ThermoFisher Scientific Kingfisher™ Flex MagMax Microbiome Ultra Nucleic Acid extraction kit and sequencing for the amplified V4 region of 16S rRNA gene took place at Argonne National Laboratory on the Illumina MiSeq platform (universal primers 515F and 806R; (35)). Illumina reads were cleaned using dada2 (36) and classified using PAthway PRediction by phylogenetIC placement (paprica v0.7.0; (37)) with a database derived reference tree (Genbank RefSeq; (38)). Point of placement was additionally used to perform metabolic inference estimating genetic characteristics, including genome size, 16S rRNA gene copy number, and GC content (39) and doubling time (gRodon; (40)). Note, doubling times from codon usage reflect theoretical growth rates, and neglect environmental conditions, serving only as a theoretical maximum to compare between samples. Additionally, taxonomic associations from paprica represent an ASV’s closest relative among the published Genbank RefSeq completed genomes and don’t always indicate the presence of an exact strain. Estimated 16S rRNA gene copy number was used to normalize read counts prior to calculations of relative abundance. Singlton ASVs were not included, and data was normalized using the Hellinger Transformation (performed in ‘vegan’; (41)), making it more suitable for downstream clustering analyses with assumptions of Euclidean distances (42).

The extraction of DNA from the 18 metagenomes is the same as described in Boulton et al. (2025) (43). Seawater was filtered through 0.22 μm Sterivex filters, and DNA was extracted using the Qiagen PowerWater DNA kit (Qiagen N.V., Hilden, Germany), following the Qiagen DNeasy Power Water SOP v1. Samples were sequenced on an Illumina NovaSeq S4 device, with 151 base pair paired-end reads, using the Illumina regular concentration library protocol. Metagenomes were assembled, annotated, and reads mapped by the JGI Metagenome Annotation Pipeline (44). To assess potential metabolic function within the 18 metagenomes, we calculated the relative abundance of COG categories (45), based on transcripts per million (tpm). The tpm abundance of each COG category was then calculated based on the normalized combined coverage of each CDS containing a COG annotation.

### Machine learning and statistical analyses

All analyses were conducted in R (46). The PCoA was constructed using the ‘vegan’ package (41) to provide a low-dimensional projection of samples based on Bray-Curtis distances which was separated into distinct clusters using the unsupervised k-means ML algorithm. The final selection of k was determined using both the elbow method, or inflection in within-clusters sum of squares (WSS), and the Silhouette method. While the elbow method identifies the point of diminishing returns in terms of cluster compactness, silhouette scores also account for between-cluster separation. Experimentally testing k values around the perceived optimums, we compared how well clusters separated in the first two ordination axes and used a post-hoc Permutational Multivariate Analysis of Variance (PERMANOVA) test to confirm that clusters were statistically significant (Fig. S1).

The unsupervised SOM model was constructed using the ‘kohonen’ package (47). Grid parameters were experimentally selected, while aiming for an optimal distribution of samples assigned to each map unit. The final map was constructed using a 4 x 4 toroidal grid with hexagonal map units and a decay scaling alpha from 0.05 to 0.01. Unit assignment was based on Euclidean distances and clustering also took place via k-means (Fig. S1). This is a similar method to what was used on a full water column MOSAiC dataset in Chamberlain et al. (2025) (15) but resulting in slightly different SOM construction and sensitivity due to the differing resolution of input data (215 underway vs. 693 samples underway and CTD samples, 0-4000 m). Resulting differences are described in detail in Figure S2. Ultimately, through the combination of dimension reduction (PCoA or SOM) and k-means clustering, each sample was assigned to unique community compositional clusters which we term as community ‘modes’. Each mode was then statistically compared to metabolic inference and environmental parameters using Kruskal Wallis and post-hoc Wilcoxon signed-rank tests (rstatix package;(48)).

The WGCNA analysis was constructed using the ‘WGCNA’ package (49) and a hybrid network construction through the application of k-means clustering instead of the standard hierarchical clustering and dynamic tree cut, inspired by the framework described in Botía et al., (2017) (50). A soft-thresholding power of 8 was chosen based on the scale-free topology fit index (R^2^ > 0.8) and mean connectivity profiles. An adjacency matrix was computed using a signed network derived from Pearson correlations, followed by transformation into a topological overlap matrix to capture shared patterns of co-abundance among ASVs. To detect modules of co-varying ASVs, we applied k-means clustering to the TOM-based dissimilarity matrix (Fig. S3). Following module detection, each module was summarized by its eigengene—the first principal component of the ASV profiles within the module. These module eigengenes were then correlated with functional metadata (estimated 16S rRNA gene copy number, doubling time, genome size, GC content, and total chlorophyll-a concentration) and environmental parameters (temperature, salinity, and location) to identify ecologically relevant modules. Module-trait relationships were visualized in a heatmap with both correlation coefficients and p-values. Similar modules were merged using a correlation-based threshold of 0.7 to reduce redundancy. To assess the importance of individual ASVs within each module, we calculated module membership scores (kME), defined as the Pearson correlation between each ASV’s abundance profile and the module eigengene. These scores were used to rank ASVs by their centrality within a module, allowing identification of the taxa with the strongest association to overall module structure and likely ecological influence.

### Data Availability

Raw data files for the 16S rRNA gene amplicon sequence data used in this study are available through the National Center for Biotechnology Information (NCBI BioProject # PRJNA895866) with metadata described through the Arctic Data Center (Chamberlain and Bowman 2022). Raw metagenomes will be made available by JGI through NCBI following PRJNA1160706 (43). Data QC and ML code pipelines are accessible upon request.

## Results and Discussion

### Initial community segmentation and environmental drivers

The underway surface water amplicon dataset contained a total of 5135 unique ASVs across 215 samples. We utilized all samples and sequences when constructing the PCoA and SOM models, which are more robust to zeros and outliers. For the WGCNA correlation network, which is sensitive to outliers and rare abundances, we reduced the training dataset to 212 samples and the most abundant 1322 sequences following standard data selection methods (49) prior to analysis. K-means clustering yielded 3 PCoA (Fig. S1), 5 SOMs (Fig. S1), and 4 WGCNA (Fig. S3) unique microbial community clusters respectively. However, SOM2 was only assigned to two samples. Both samples were from a period of lower latitude transit through the marginal ice zone and likely represented a brief spatial disruption in community succession. This is reflected in both the PCoA, which clustered these samples with the later dominant PCoA3, and WGCNA, which showed a brief inversion in the dominance of the Blue and Brown modules at this time (Fig. 1). These two samples are either true outliers in the dataset, and the SOM map grid was the most sensitive projection, or it was a cut-off induced by insufficient grid-size or misinterpretation of map proximity (22). Because K-means clustering assumes clusters of equal variance, non-balanced clusters can result in unduly isolated, low-variance clusters (51). While SOM2 and SOM4 (nearest neighbor, Fig. 1d) were not significantly different in most genetic parameters, there was some significant environmental variation and SOM2 did display a significantly higher GC content (p = 0.0062; Fig. S5). However, with only n = 2, we were not confident in these relationships and SOM2 was not considered functionally unique from SOM4 in further downstream analyses and comparisons.

**Figure 1.**
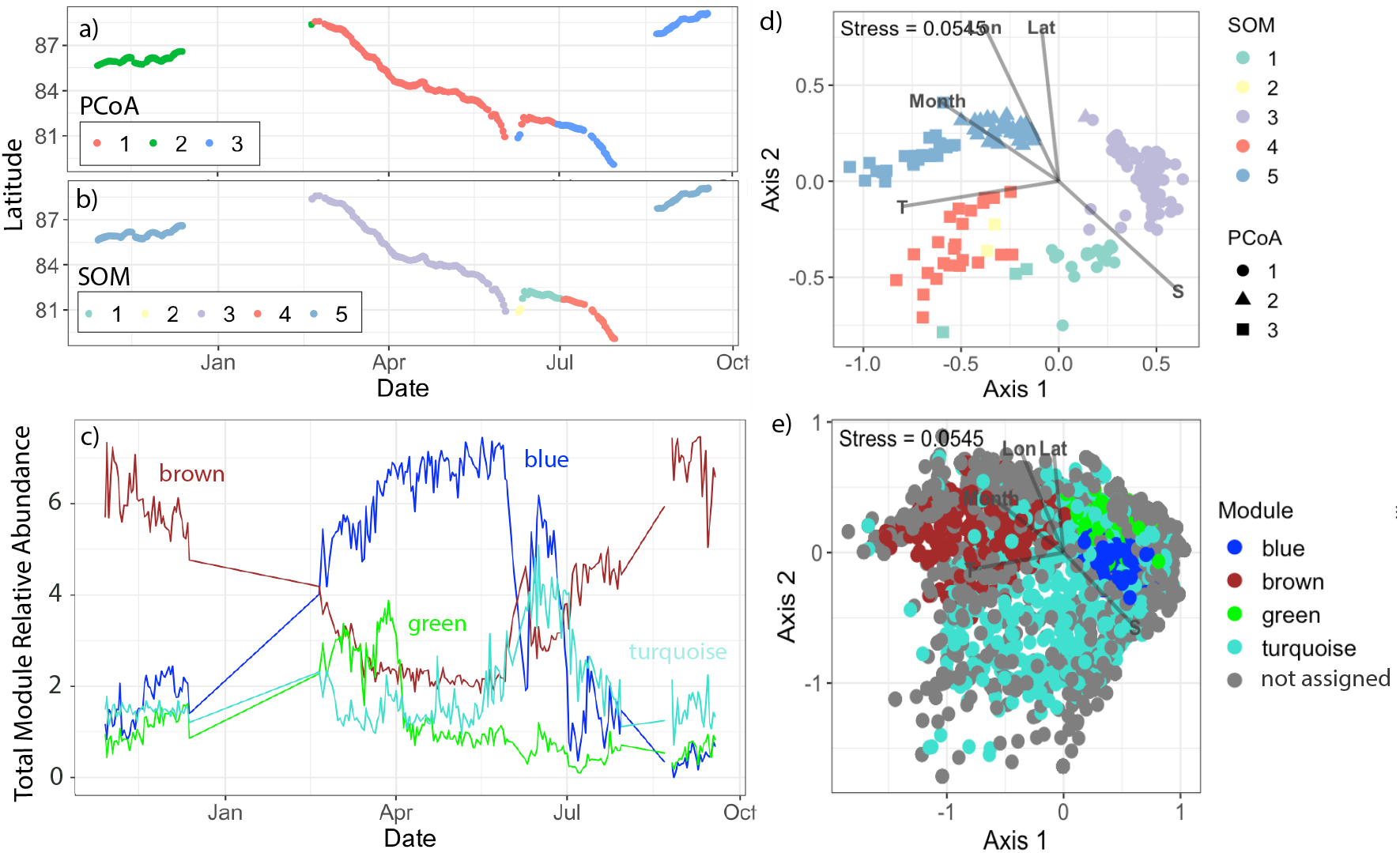
Clustering output and spatio-temporal distributions during MOSAiC. PCoA (a) and SOM (b) clusters identified distinct seasonal shifts in bacterial community structure throughout the MOSAiC Drift, here visualized as latitude. In (c), the relative abundances of all ASVs identified to each WGCNA module were summed for each sampling date. Environmental separation among clusters was significant (indicated by the grey arrows for latitude (Lat), longitude (Lon), numerical month, temperature (T), and salinity (S)) when samples (d) and ASVs (e) were visualized in the first two axes of an NMDS ordination created from the bray dissimilarities of Hellinger transformed 16S ASV relative abundances. The NMDS stress value, indicating ordination fit, is listed in the top left-hand corner. In (d) samples are colored by SOM and shaped by PCoA cluster assignment. In (e) ASVs are colored by module assignment. Grey points represent ASVs with total relative abundances too low to be included in the WGCNA analysis.

Overarching seasonal clusters remained generally the same across models, but there was substantial variance between the temporal cut-offs for each methodology (Fig. 1). When projecting the underlying data structure of both the samples and individual ASVs using an alternative projection method (NMDS ordination) (Fig 1d, e), separation in the ordination space was most closely aligned with the SOM segmentation (samples) and WGCNA modules (ASVs). Numerical month (r^2^ = 0.56), temperature (r^2^ = 0.66), salinity (r^2^ = 0.70), latitude (r^2^ = 0.65), and longitude (r^2^ = 0.79) were all significant explanatory variables (p < 0.001) in the NMDS ordination (Fig 1d, 1e). The strong drivers of longitude and salinity are notable, but not unexpected. Longitude (and latitude) represent the large-scale oceanographic gradients shaping community composition. Salinity (and temperature) are related to these gradients but also impacted by seasonal ice melt – which created critical thresholds for community structure and ML clustering (Table 1). The PCoA was the least sensitive to these finer environmental shifts, recognizing only one community cluster both during summer melt and in fall 2020 (Fig. 1), when MOSAiC drifted into relatively fresher Transpolar Drift influenced waters (33). Meanwhile, the period between June–September (summer melt season) had three distinct co-occurrence modules (brown, turquoise, and blue; Fig. 1c) and two unique SOM communities (SOM1 and SOM4; Fig. 1b), indicating greater sensitivity of these methods to recognizing fine scale environmental and/or seasonal shifts in community structure. This advantage likely stems from the underlying data structure. PCoA axes are continuous and dominated by highly abundant taxa (42). Rare taxa get compressed near the origin within the ordination, meaning their influence is likely to be under-represented (52). While this projection is useful in analyses specifically targeting a balanced representation of the community, our results indicate this may come at the cost of functional sensitivity and support other studies which have highlighted a mismatch between abundance and influence (9).

**Table 1.** Functional summary of surface ocean Arctic Ecotypes.

| Description | Growth rate estimate | Estimated genome size | Key taxonomic representatives | Enriched cellular functions | ML clustered output representatives |
| --- | --- | --- | --- | --- | --- |
| Bloom Associated | Fastest | Large | Polaribacter, Sulfitobacter | Motility and cell division | SOM1, Turquoise module |
| Post-Bloom | Fast | Large | Flavobacteria, general | Carbohydrate and lipid processing | PCoA3, SOM5, Brown module |
| Winter/Dark | Slowest | Streamlined | Various Gammaproteobacteria | Energy production and conversion | Green module |
| Spring/Light | Slow | Streamlined | Nitrospulimus sp., Pelagibacter, Phycosp | Defense mechanisms, inorganic ion transport | PCoA2, SOM3, Blue module |

Annually recurrent microbial ecotypes have been observed in other Arctic Ocean time series (27), due to strong seasonal selection pressures (53). The SOMs effectively captured this seasonal succession because the entire community assemblage shifted coherently. The ASV-level WGCNA modules additionally emphasized background inter-seasonal aspects of the dataset (Green module). By building a signed correlation network among individual ASVs that co-vary across samples, the modules emerge from shared temporal dynamics, not simply relative abundance. This explains why the method was more sensitive to dynamic environmental shifts – allowing it to better resolve fine-scale resource niches (Turquoise module).

### From microbial diversity to microbial function

Seasonal cutoffs between most clusters were reinforced by clear taxonomic and genetic shifts. SOM5, for example, is present in both late Fall 2019 and early Fall 2020 and was enriched in ASVs assigned to taxa in the Brown module, including some of the top module members (e.g. *Planktomarina temperata*, and an unidentified Flavobacterium ASV; Fig. 2). Most of these top taxa represent copiotrophs associated with the rapid colonization and degradation of organic material (54). SOM5 samples were additionally enriched in carbohydrate, amino acid, and lipid transport genes (Fig. 3). The PCoA analysis however, distinguished two clusters during this same time (PCoA2 in Fall 2019 and PCoA3 in late Summer/early Fall 2020), driven by overall water mass properties of salinity and temperature (Fig. 1d). The primary taxonomic difference between these modes were the additional enrichment in ASVs assigned to *Bermanella marisrubi, Owenweeksia hongkongenesis* and *Polaribacter* sp. (Fig. 2) and cell motility/translational genes (Fig. 3) in PCoA3. However, these features are more prominent in the additional SOM cluster of SOM4 which is only present in July 2020 and additionally marked by significantly lower mean doubling times (Fig. 4), higher mean 16S copy numbers, and higher mean genome sizes (Fig. S5). While the July 1 shift between PCoA2 and 3 coincide with the onset of Brown module dominance, the taxa with both high kME and significant negative correlation to mean doubling (Fig. 4) align exactly with the topmost abundant taxa in SOM4. This alignment indicates that these taxa are their own unique, and fast-growing community which could functionally be recognized as its own group.

**Figure 2.**
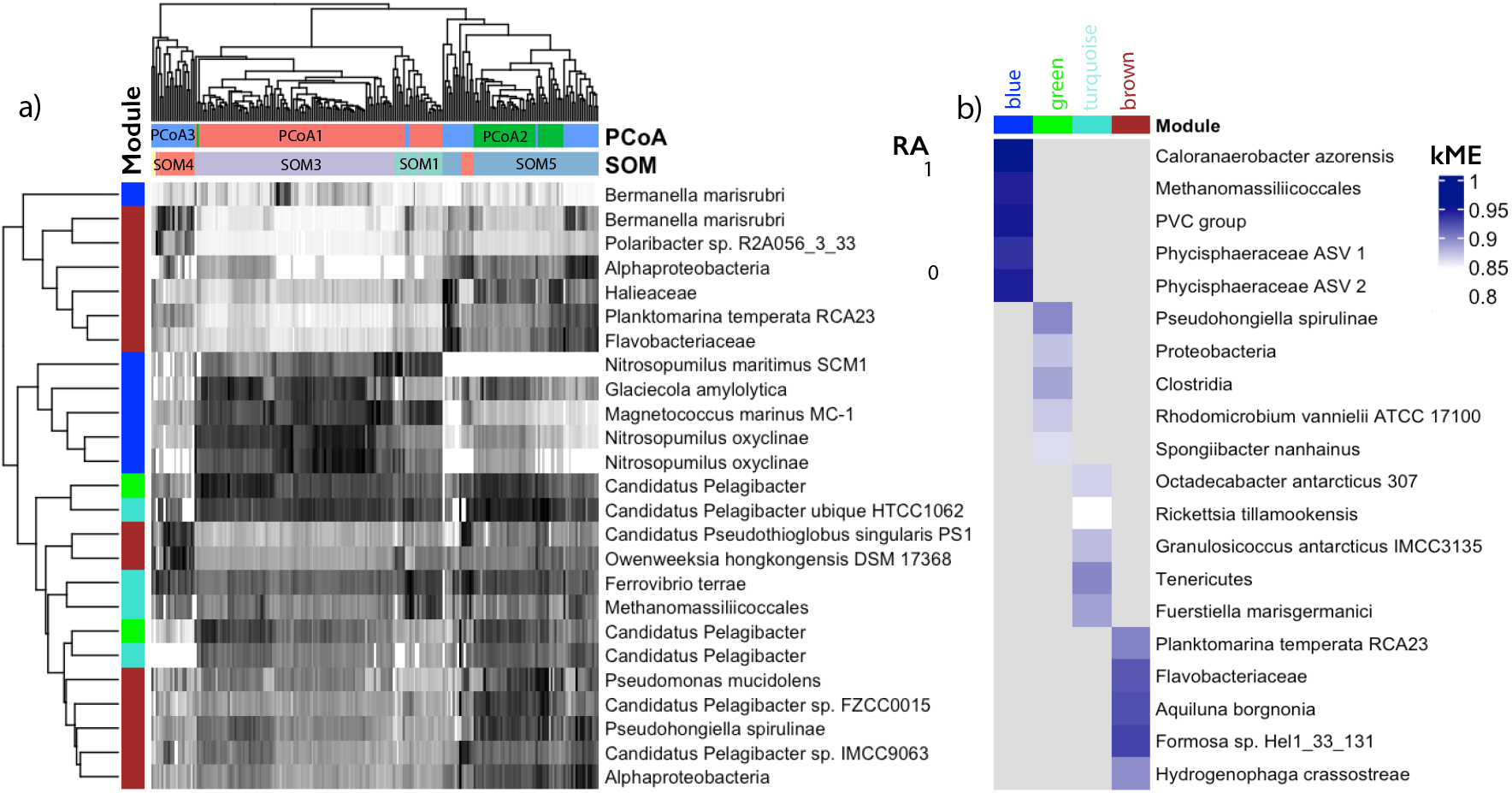
Abundant taxonomy of ML clusters and top WGCNA module membership. (a) Heatmap of Hellinger transformed relative abundance (RA, scaled 0-1 via z-score) for the 25 most abundant ASVs across all samples. Columns are annotated by PCoA and SOM cluster assignments, and rows are colored by WGCNA module membership. Both were clustered using euclidean distances. (b) Heatmap showing the ASVs with highest module membership (*k*ME) for each WGCNA module, with color labels matching the rows in panel (a).

**Figure 3.**
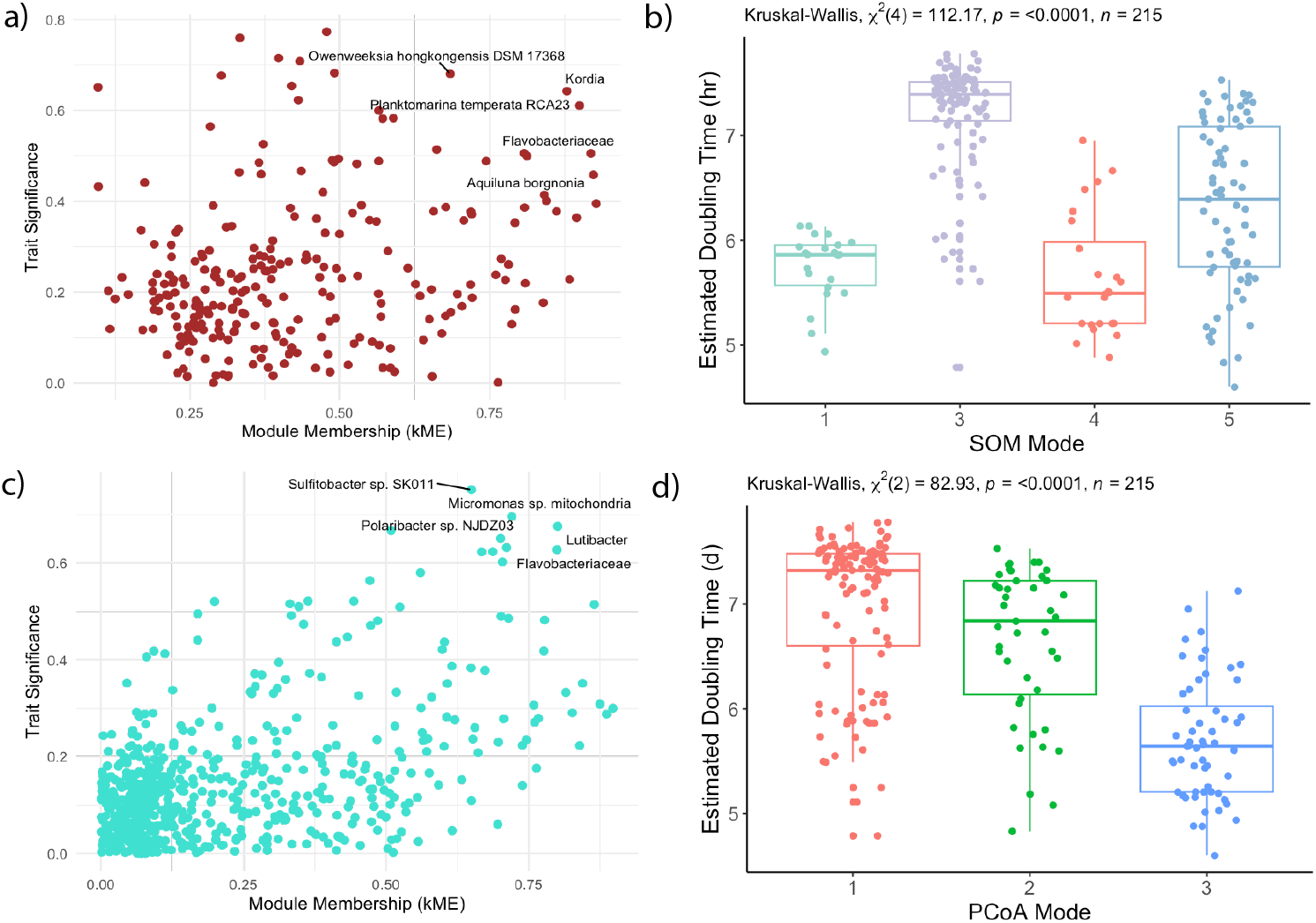
Clusters of Orthologous Groups (COG) enrichment across sample clusters. Top 25 most abundant COG (Clusters of Orthologous Groups) assignments using Pfam abundance (tpm) across all samples. Columns represent the 15 sample days where surface water metagenomic analyses aligned with underway 16S sampling. Columns are color annotated by PCoA and SOM cluster assignments as well as the dominant WGCNA module for that date. Rows correspond to COG assignments with 4 containing functional genes enriched in multiple COG groups simultaneously. Abundance values were normalized (tpm) and scaled from 0-1 using z-score calculations for plotting. Hierarchical clustering was applied to both rows and columns.

**Figure 4.**
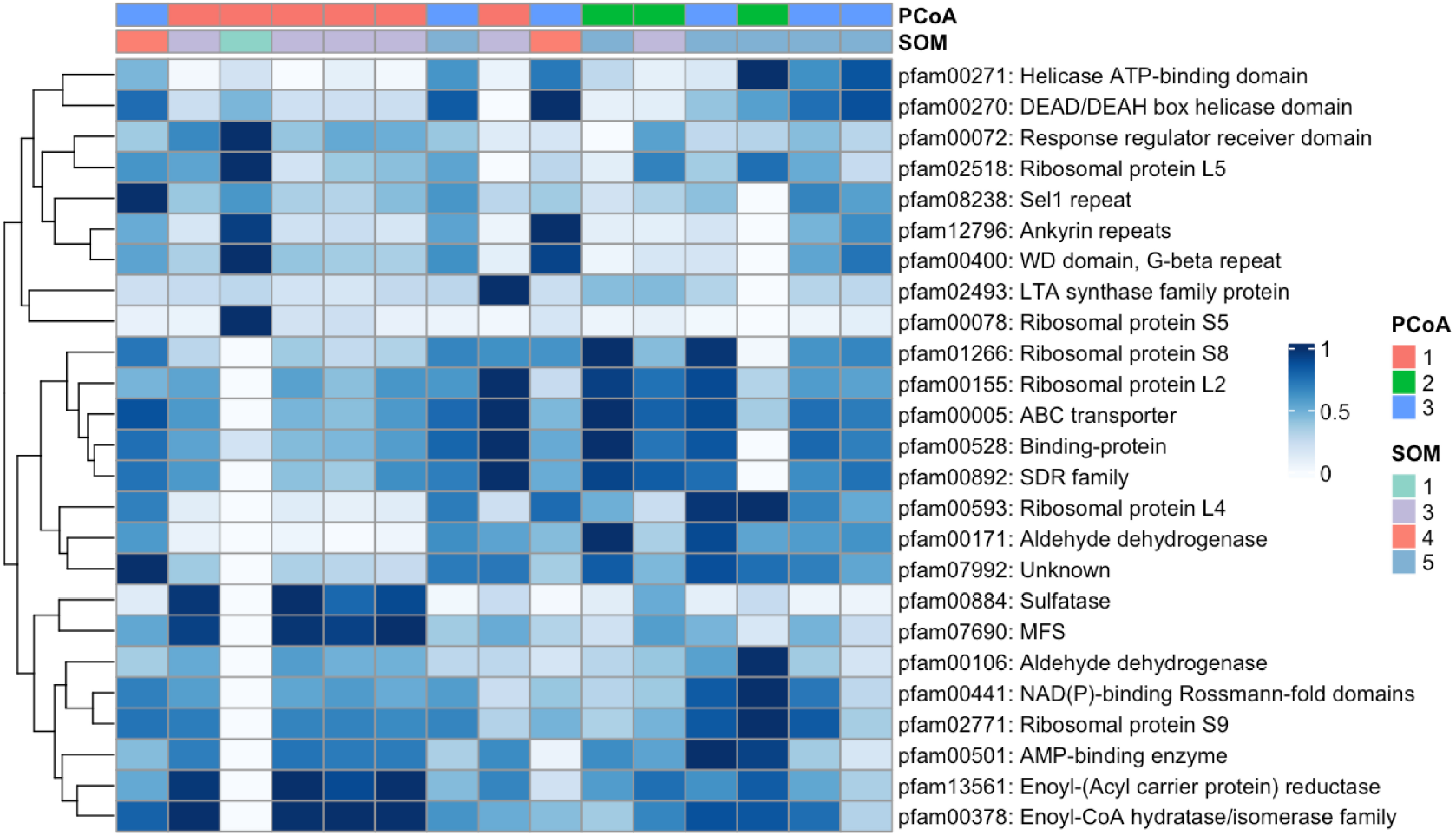
Codon-usage derived growth rates across clusters and highly correlated taxa. Growth rate is represented here by the gRodon estimated mean doubling time in hr, where a lower doubling time indicates faster growth. (a) Scatterplot of trait significance (doubling time) and module membership (kME) for all ASVs in the Brown module. (b) SOM clusters compared to doubling time. (c) Scatterplot of trait significance (doubling time) and module membership (kME) for all ASVs in the Turquoise module (d) PCoA clusters compared to doubling time. In panels (a) and (c), the top 5 significantly correlated and high kME ASVs are labeled by closest reference genome. In panels (b) and (d), the horizontal line of each boxplot indicates the median, the box spans the first to the third quartile, the whiskers indicate minimum and maximum values, and the dots indicate all values

Despite being the module with the most ubiquitous background taxa (Fig. 1e), the Turquoise module was significantly (p < 0.001) negatively correlated with mean doubling time, with its top kME and correlated taxa strongly related to phytoplankton groups (e.g. *Micromonas* mitochondrial DNA) or associated heterotrophs known for exploiting sudden pulses of organic matter (e.g. Polaribacter, Lutibacter). The Turquoise module was also the most significantly correlated with chlorophyll-a concentration (p < 0.001; Fig. S6) and had several high kME ASVs assigned to either phychrophilic biofilm producers or algal associates/pathogens. Its sudden increase in abundance during the period of the pelagic phytoplankton bloom (mid-late June;(31)), indicates a shift in carbon source utilization and increased growth rates (high enrichment of cell division genes, Fig. 3). This resulted in a unique bloom-associated community which grew in abundance during this time. These functional changes in genetic information were captured with the distinct segmentation of SOM1, had high chlorophyll-a concentrations, the second lowest mean doubling times, second highest mean genome size, and a low 16S copy number (Fig. S5). However, this SOM cluster was also composed of highly abundant ASVs assigned across the Blue, Brown, and Turquoise modules, which were all relatively abundant during this seasonal transition as well (Fig. 1). While there was only one matching metagenome sample for this period, it was enriched in unique genes related to the mobilome, cell division, and transcription (Fig. 3) supporting distinct functionalities and rates during this time (Table 1).

The only true inter-seasonal community dynamics masked by the SOM segmentation was the appearance of the Green module in early spring, prior return of sunlight at the end of March. Its most abundant ASVs belonged to ASVs assigned to the slow growing and genomically streamlined Pelagibacterales order ((55); Fig. 2a), which were also highly abundant in the Blue module and are common for the Arctic winter and spring in the upper water column (15). Pelagibacterales are also uniquely distinguished by their high internal diversity – even within subclades, the micro-diversity of this species is extremely high, allowing them to track subtle shifts in environmental niches across the world’s oceans (56). It is therefore not surprising that they are highly abundant across multiple ML clusters as there is likely greater speciation not captured in the reference genomes used here for taxonomic assignment. The ability to capture meaningful ecological differences and functional separation even without perfect taxonomic assignment highlights one of the primary benefits to using ML segmentation at the ASV-level.

While the broader winter/spring trends were captured by PCoA1 and SOM3 (Fig. 1), it is likely that Green module taxa are responsible for the large spread of doubling time estimates in both these sample-driven clusters (Fig. 4). Generally, however the Green module followed the same trait-relationships as the more abundant Blue module (Fig. S6). The only exception is an opposite correlation to Latitude, potentially indicating oceanographic partitioning at this time, although the decrease in Green module relative abundance occurs slightly after the Polarstern drifted across the Gakkel Ridge (33). Additionally, the top kME taxa for this module (Fig. 2b) are primarily specialized in breaking down recalcitrant carbon and their genomic representatives all contain at least one mechanism for boosting ATP yields during periods of electron scarcity (i.e., rhodopsins and other bacterial photosystems, motility). This indicates their suitability for surviving the scarcity period of polar night and adapting early to the return of the sun in early spring, as seen in the overwintering strategies of Arctic phytoplankton and sea-ice algae (57, 58). The Blue module on the other hand is driven by phytoplankton associated and chemosynthetic taxa (Fig 2b), which were brought to the surface from deeper in the water column due to intensified mixing at this time (15) and representing a total functional shift that is best reflected in the most abundant ASVs in SOM3 (Fig 2a). SOM3 is also enriched in heterotrophic metabolism gene indicators, as well as defense mechanism genes – which would reflect the growing number of phytoplankton as the Spring and potentially grazing activities by mixotrophic dinoflagellates, which were also brought to the surface and highly abundant at this time (15).

The functional organizations offered by ML methods (Table 1) may inform ongoing debates over microbial functional redundancy, a concept used to explain the stability of many microbial processes despite high individual species turnover (59) The distinct temporal windows occupied by the SOM clusters and WGCNA modules and clear shifts in community composition coinciding to measurable changes in dominant metabolic function (Fig. 3), support studies emphasizing a tight coupling between taxa and function (60) and limited redundancy at ecologically relevant scales of environmental change (61). As has been found in other study systems (62), these two methods revealed a layered picture of community succession, where seasonal shifts were functionally buffered (SOMs/PCoA) but inter-seasonal responses to substrate pulses depended on specific, and not fully redundant, functional guilds that the biologically informed co-occurrence WGCNA analysis was able to extract.

### Implications for integrating ML with mechanistic biogeochemical models

A central challenge in ocean modeling is representing the immense diversity of marine bacteria. Most process-based biogeochemical models either do not explicitly represent bacteria (63) or rely on overly simplistic approaches, such as a single bacterial compartment that fails to capture diverse metabolic strategies and environmental responses(64, 65). Although this is beginning to change as new studies address this shortcoming (e.g., Hasumi & Nagata, 2014 (66); Kim et al., 2023 (67); Lennartz et al., 2024 (68); Zakem et al., 2025 (69)), the resulting mismatch between complex bacterial processes and their simplified representations undermines our ability to accurately quantify the bacterial role in climate and carbon cycle dynamics.

Our results demonstrate that ML approaches provide an efficient solution to the challenge of scaling between individual taxa and bulk ecosystem compartments by creating functionally meaningful intermediate units of complexity. We identified four distinct ecotypes in the surface Arctic Ocean, each characterized by clear environmental drivers and seasonal patterns. Two ecotypes dominated during winter and spring periods characterized by low light conditions, while two others were tightly coupled to the timing of summer phytoplankton blooms (Table 1). These ML-derived ecotypes represent ecologically interpretable functional units ideal for model integration because they consolidate key physiological attributes including growth strategies (16S copy number), metabolic potential (COGs), and salinity tolerance. Moreover, these approaches generate ready-made state variables whose parameters, such as maximum growth rates, temperature-salinity optima, and substrate preferences, can be constrained from multi-omic datasets, eliminating dependence on outdated single-species culture estimates. Recent studies have demonstrated significant improvements achievable through incorporating such bacterial trait structures (69).

We propose treating each ML ecotype as an emergent functional type with parameter values derived from ensemble trait measurements (e.g., median Vmax, half-saturation constants). Because these ecotypes maintain clear relationships with environmental covariates such as salinity, temperature, and longitude, their abundances can be dynamically adjusted using environment-dependent growth and loss terms. This approach adds adaptive realism to models without increasing computational complexity. In doing so, the selection of appropriate ML methods is critical. The clear difference between our ordination approaches highlights the necessity of employing non-linear, dimensionality-reduction techniques when deriving model compartments. While traditional axis-based ordinations (PCoA) grouped samples solely by water mass characteristics, they failed to capture the seasonally coupled salinity-temperature niches that more sophisticated methods like SOM and WGCNA successfully identified. Integrating comparable ML segmentation within model-data assimilation frameworks could substantially improve state estimation and parameter optimization (70, 71), ultimately strengthening constraints on bacterial feedback to ocean carbon cycling.

## Conclusions

ML provides a workable bridge between microbial surveys and the coarse compartments required by biogeochemical modeling. Our results indicated that functional variation and shifts in species interaction were accurately captured by ML segmentation – most notably SOMs – and we were able to identify “functionally coherent” groupings which crossed taxonomic boundaries (e.g. Pelagibacterales, Flavobacteria) and embodied distinct variations in the genetic traits which are most useful for predictive model parameterization. The utility of compartmentalizing the community in this way is that it goes beyond ‘size-class’ based trait extraction under the argument of functional redundancy and still allows groupings when taxonomic succession does matter. Furthermore, testing our workflow on amplicon sequencing derived ASVs demonstrates how these algorithms could be used to combine and mine previously generated legacy data for new ecological insight. Amplicon sequencing has long been the more widespread and affordable methodology compared to the richer but scarcer metagenomic and transcriptomic records. However, this work could still be improved – particularly in the Arctic where data is scarce – by producing more *paired* datasets for such ML methods to get better predictive insights into the mechanisms behind biogeochemical parameters of interest in a rapidly changing ocean.

## Acknowledgements

This is the National Science Foundation (NSF) Center for Chemical Currencies of a Microbial Planet (C-CoMP) publication #77. Amplicon sequence data collection and processing for MOSAiC was funded by the US NSF award, OPP 1821911, to J.B. This award also supported MOSAiC participation for J.M.C, J.B. and E.J.C. J.M.C was additionally supported by the US Department of Energy (DOE) Atmospheric System Research program (grant DE-SC0022046) and E.J.C was additionally supported by an NSF Graduate Research Fellowship. W.B. was supported by the Natural Environment Research Council and ARIES DTP (grant NE/S007334/1). Metagenome sequencing (project 10.46936/10.25585/60001271) was carried out by the DOE Joint Genome Institute (JGI) in a User Facility supported by the DOE Office of Science (grant DE-AC02-05CH11231). Development of this manuscript and T.C. was supported by an NSF Postdoctoral Research Fellowship award, OPP 2317681, to E.J.C. HHK was supported by C-CoMP (NSF Award OCE-2019589).

The data used in this research was collected as part of the Multidsiciplinary Drifting Observatory for the Study of Arctic Climate (MOSAiC) project (data tag MOSAiC20192020) and we acknowledge and thank all persons involved in this expedition as per the MOSAiC extended acknowledgement (72). We would especially like to thank all members of Team Ecosystem (31) and specifically the Eco-Omics team (10), led by Katja Metfies and Thomas Mock.

Author CRediT: Emelia J. Chamberlain conceptualized the study and wrote the original draft of the manuscript. Jessie M. Creamean, Jeff Bowman, and Emelia J. Chamberlain collected samples in the field. Jeff Bowman and Emelia J. Chamberlain performed the amplicon data curation. William Boulton and Thomas Mock conducted the metagenome data curation. Emelia J. Chamberlain performed the formal analysis and visualization with assistance from Theodore Calianos and input from Elizabeth Connors. Heather H. Kim supervised the project and edited the original draft of the manuscript. All authors contributed to manuscript review and approved the final version for publication.

## Supplemental Material

**Figure S1.**
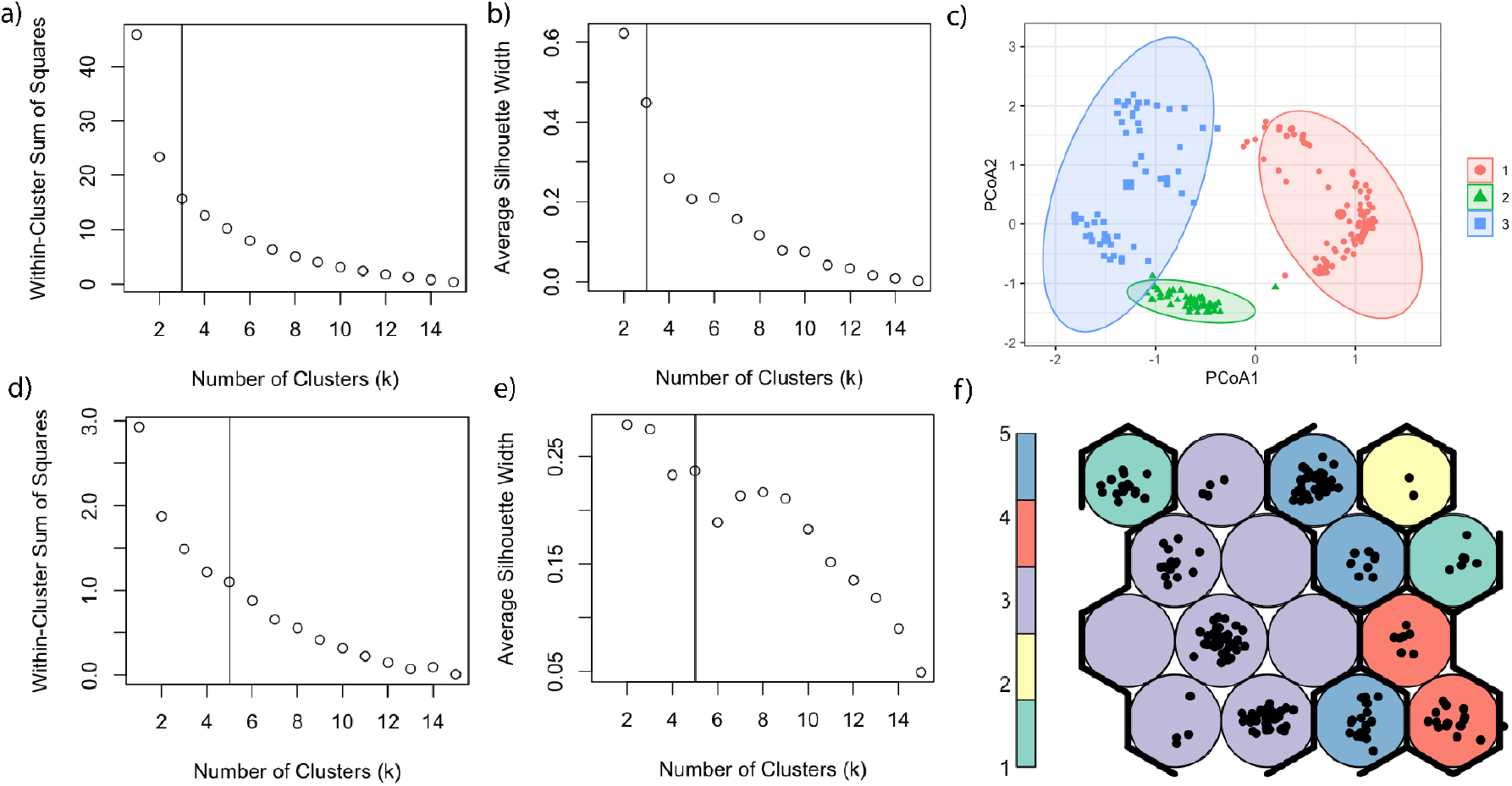
Comparison of PcOA and SOM projections with k-means clustering (samples). The within-cluster sum of squares (a,d) and average silhouette width across increasing *k* values (b,e) for clustering in PCoA space (a,b) and across SOM vectors (d,e). The PCoA ordination (c) shows samples colored and shaped by k-means cluster assignment (k = 3), with 95% confidence ellipses. SOM grid (f) shows map nodes colored by final k-means cluster assignment (k = 5), with black dots representing individual samples.

**Figure S2.**
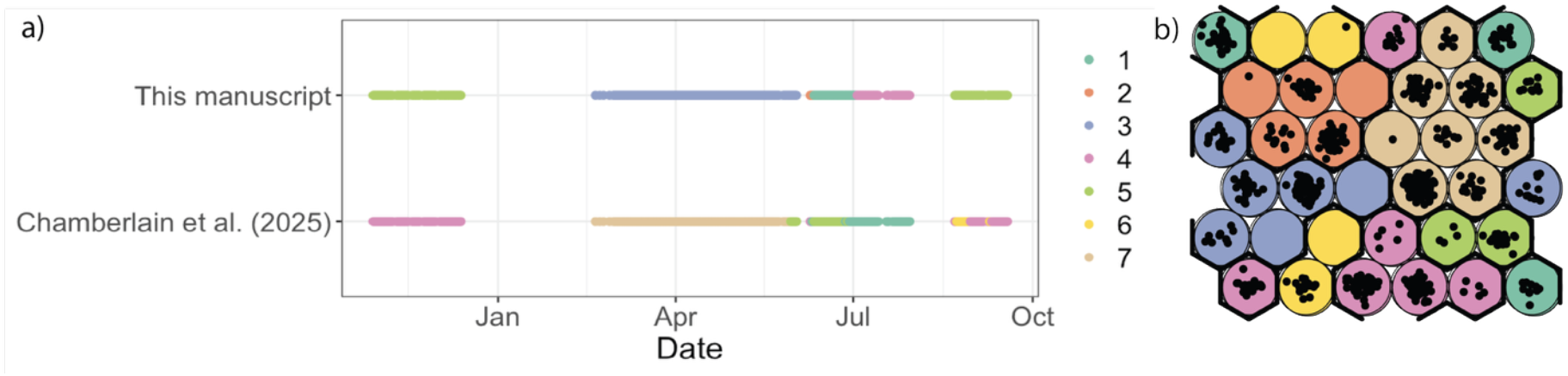
Comparison of MOSAiC SOMS constructed using only underway data (Fig. S1) and full water column data (Chamberlain et al. 2025). All underway DNA samples are presented across the MOSAiC time series (a) colored first by the SOM mode assignments trained using only 11 m underway samples in this manuscript (Fig. S1f), and then by SOM mode assignments trained using a 693 sample full water column dataset including both 11 m underway and 2–4000m CTD water samples (Chamberlain et al. 2025, panel b). The incorporation of additional and shallower surface water samples from the CTD casts results in a more sensitive model and an additional community mode representative of river-influenced water from the Transpolar Drift (Chamberlain et al. 2025). Seasonal modes and their transitions remain otherwise nearly identical between the two model constructions. Due to the greater information provided for training, the Chamberlain et al. (2025) is likely the more accurate representation of community composition, and we recommend using this model in future studies beyond this methodological comparison.

**Figure S3.**
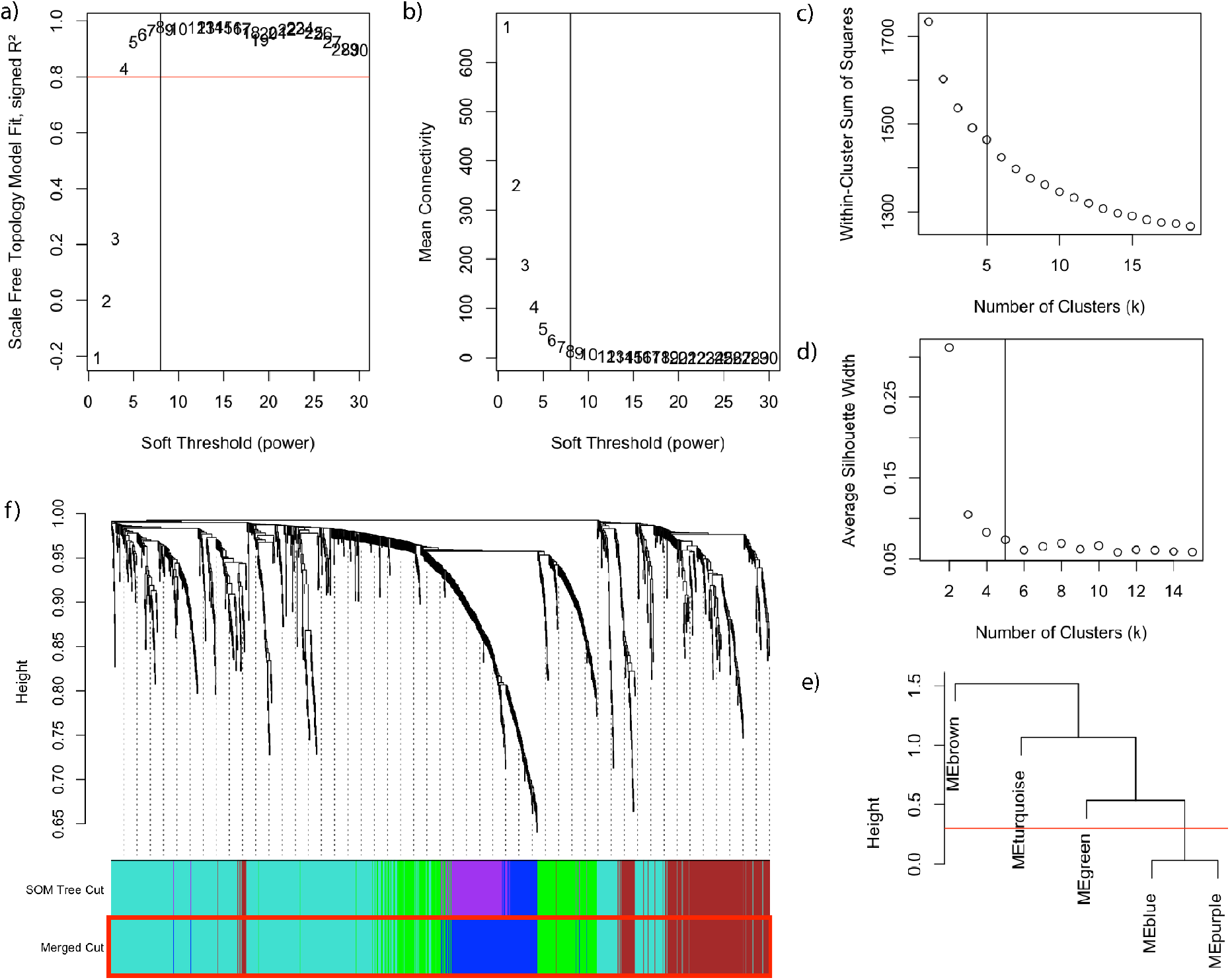
WGCNA clustering (ASVs) using k-means for module detection. Panel (a) shows the scale-free topology model fit (signed R^2^) plotted against soft-thresholding powers, where the red line indicates a threshold of R^2^ = 0.8. Panel (b) highlights the mean connectivity across the same range of powers. Both a scree plot of within-cluster sum of squares (c) and the average silhouette width (e) were used to determine the optimal number of module clusters (k = 5). To confirm our selection, a clustering dendrogram of the module eigengenes was constructed and branches with high similarity (< 0.3, red line) were merged (final k = 4). The dendrogram of included ASVs (f) indicates module assignment, showing both the original SOM-based module cut and the final selection of merged modules (red box).

**Figure S4.**
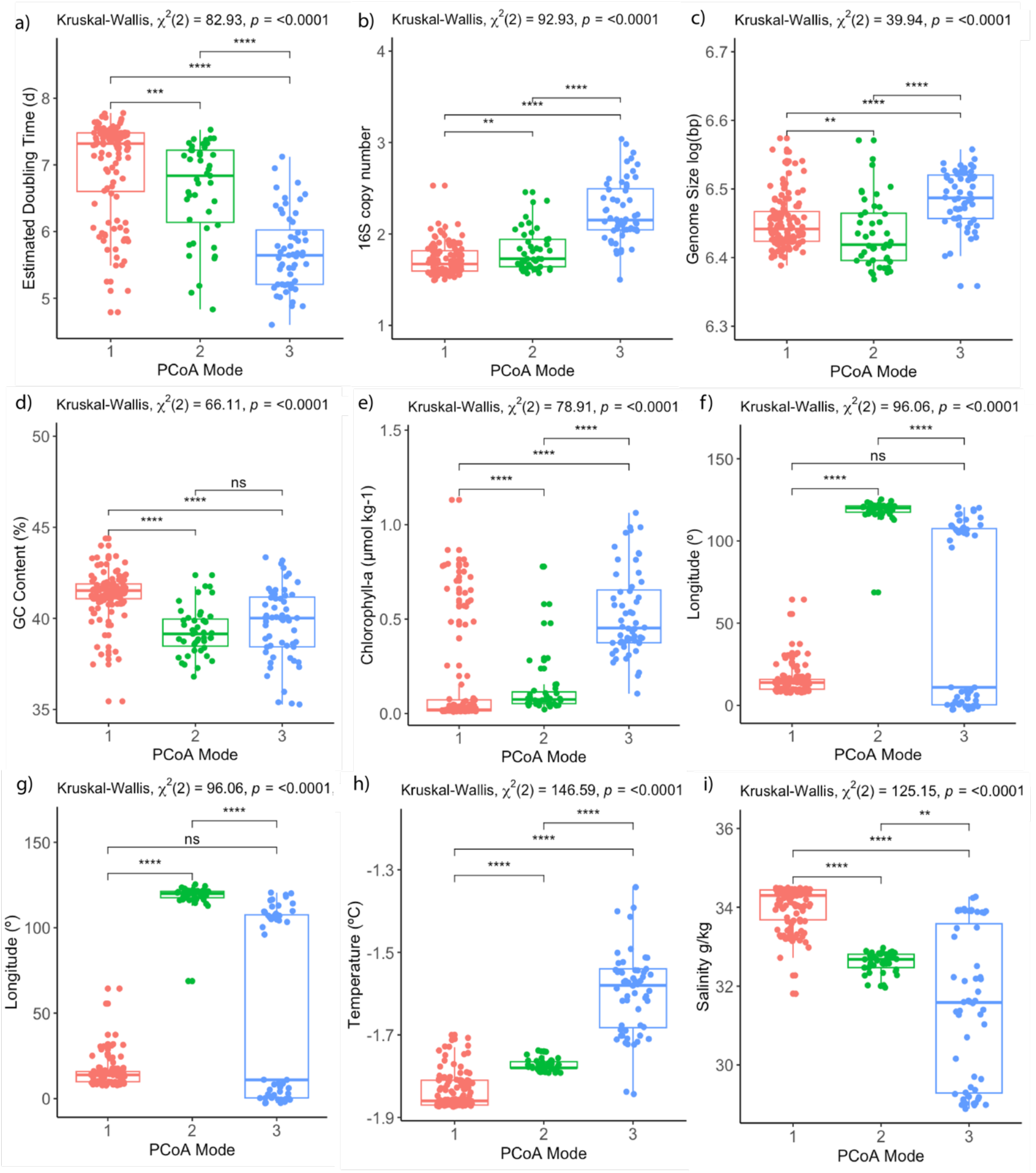
PCoA distribution across genomic and environmental parameters. Where paired samples were available, SOM cluster is compared to a) gRodon estimated doubling time in hours, b) estimated 16S rRNA gene copy number, c) estimated genome size in bp, logged, d) estimated % GC content, e) chlorophyll-a concentration in µmol kg^-1^, f) latitude in º, g) longitude in º, h) temperature in ºC, and i) salinity in g kg^-1^. The horizontal line indicates the median, the box spans the first to the third quartile, the whiskers indicate minimum and maximum values, and the dots indicate all values. The results of a Kruskal-Wallis and post-hoc Wilcoxon-signed rank test comparisons are presented in brackets where ns indicates p > 0.05, * indicates 0.01 < p ≤ 0.05, ** indicates 0.001 < p ≤ 0.01, *** indicates 0.0001 < p ≤ 0.001, and **** indicates p ≤ 0.0001.

**Figure S5.**
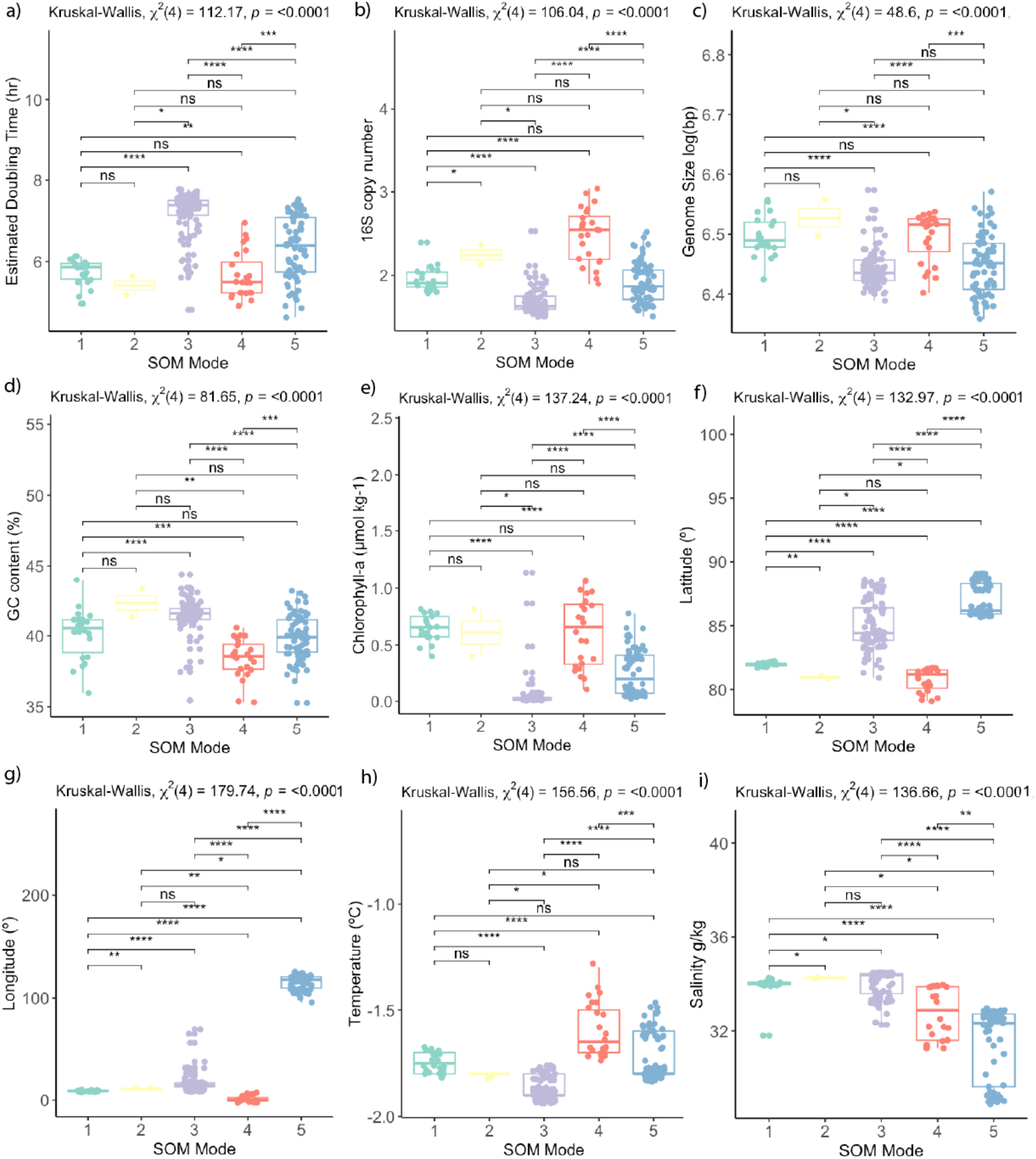
SOM distribution across genomic and environmental parameters. Where paired samples were available, SOM cluster is compared to a) gRodon estimated doubling time in hours, b) estimated 16S rRNA gene copy number, c) estimated genome size in bp, logged, d) estimated % GC content, e) chlorophyll-a concentration in µmol kg^-1^, f) latitude in º, g) longitude in º, h) temperature in ºC, and i) salinity in g kg^-1^. The horizontal line indicates the median, the box spans the first to the third quartile, the whiskers indicate minimum and maximum values, and the dots indicate all values. The results of a Kruskal-Wallis and post-hoc Wilcoxon-signed rank test comparisons are presented in brackets where ns indicates p > 0.05, * indicates 0.01 < p ≤ 0.05, ** indicates 0.001 < p ≤ 0.01, *** indicates 0.0001 < p ≤ 0.001, and **** indicates p ≤ 0.0001.

**Figure S6.**
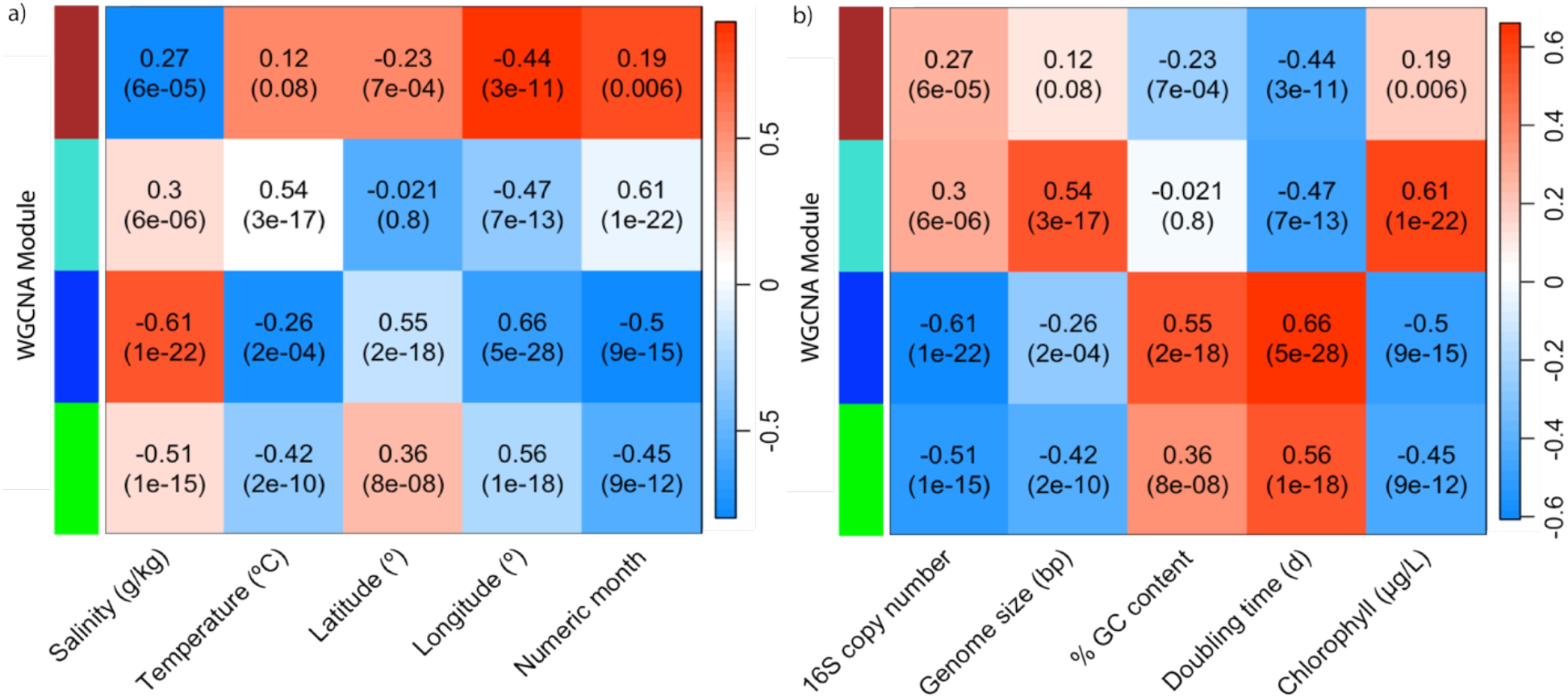
WGCNA module correlations to genomic and environmental parameters. The Pearson correlation coefficients between module eigengenes and environmental variables (a) or genetic traits (b) are shown as heatmaps. The text in each cell displays the correlation and associated *p*-value and cells are colored by the strength of their correlation (blues indicate negatively correlated relationships while reds indicated positively correlated relationships). Row colors correspond to the WGCNA module assignments of Brown, Blue, Green, and Turquoise.

